# Macromolecular Regulators Have Matching Effects on the Phase Equilibrium and Interfacial Tension of Biomolecular Condensates

**DOI:** 10.1101/2021.02.18.431854

**Authors:** Konstantinos Mazarakos, Huan-Xiang Zhou

## Abstract

The interfacial tension of phase-separated biomolecular condensates affects their fusion and multiphase organization, and yet how this important property depends on the composition and interactions of the constituent macromolecules is poorly understood. Here we use molecular dynamics simulations to determine the interfacial tension and phase equilibrium of model condensate-forming systems. The model systems consist of binary mixtures of Lennard-Jones particles or chains of such particles. We refer to the two components as drivers and regulators; the former has stronger self-interactions and hence a higher critical temperature (*T*_c_) for phase separation. In previous work, we have shown that, depending on the relative strengths of driver-regulator interactions and driver-driver interactions, regulators can either promote or suppress phase separation (i.e., increase or decrease *T*_c_). Here we find that the effects of regulators on *T*_c_ quantitatively match the effects on interfacial tension (*γ*). This important finding means that, when a condensate-forming system experiences a change in macromolecular composition or a change in intermolecular interactions (e.g., by mutation or posttranslational modification, or by variation in solvent conditions such as temperature, pH, or salt), the resulting change in *T*_c_ can be used to predict the change in *γ* and vice versa. We also report initial results showing that disparity in intermolecular interactions drives multiphase coexistence. These findings provide much needed guidance for understanding how biomolecular condensates mediate cellular functions.

## INTRODUCTION

Biomolecular condensates formed via phase separation often appear as micro-sized liquid droplets. The surface tension, *γ*, at the interface between the dense phase and the surrounding bulk phase is a very important property that affects the dynamics, organization, and, ultimately, function of biomolecular condensates. Notably, interfacial tension drives the fusion of droplets and is thus a main determinant of fusion speed.^1-3^ Experimental studies have revealed that, instead of a single homogeneous phase, condensates, including membraneless organelles such as nucleolus, can occur as multiple, coexisting dense phases, each with a distinct composition.^1, 4-10^ Interfacial tension may play a vital role in the spatial organization of the multiple dense phases.^1^ Moreover, the material states of condensates often evolve over time, both for normal cellular functions and as an aberrant transitions.^11^ The time evolution is also partly driven by material properties including interfacial tension.

Inside cells, condensates often comprise dozens to hundreds of macromolecular components.^4^ The complex composition provides ample opportunities for the perturbation of interfacial tension, by changes in macromolecular composition or their interactions through mutations, posttranslational modifications, and variations in solvent conditions such as temperature, pH, and salt. It is very possible that compositions of condensates are tailored to achieve desired interfacial tensions.^1^ However, there is no general understanding of how interfacial tension depends on macromolecular composition.

Many theoretical and computational studies have focused on calculating binodals, i.e., coexistence curves between dense and bulk phases.^12-20^ These studies, even when based on highly simplified models, have generated far-reaching conclusions on phase equilibrium. In particular, results for spherical particles and polymer chains, as models for structured and disordered proteins, respectively, have shown that chain systems tend to have a higher critical temperature (*T*_c_) and lower densities in both phases.^14^ Most relevant for the present study, our computations led to general predictions for how *T*_c_ is perturbed by compositional changes.^15, 18^ These predictions were validated by our own experiments and also explain many experimental observations reported in the literature.^18^ Extending such computations to predict interfacial tension was the main motivation for the present work.

The calculation of interfacial tensions for phase-separated model systems has a long history.^21-22^ Here we note only a few studies that have direct relevance to the present work. Lee et al.^23^ and Stephan et al.^24^ both reported interfacial tensions for binary mixtures of Lennard-Jones particles. The latter study presented interfacial tensions when the strength of the cross-species interactions was varied. Silmore at al.^25^ reported interfacial tensions for pure Lennard-Jones chains at different lengths. Our work here covers binary mixtures of both Lennard-Jones particles and Lennard-Jones chains, over a range of temperatures. Our focus is the relation between *T*_c_ and *γ* at a given temperature when the molar ratio of the two species is varied.

The Lennard-Jones interaction energy between two particles is

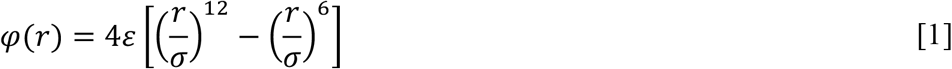

where *r* is the interparticle distance, *ε* represents the strength of the attraction, and σ represents the diameter of the particles. For a pure Lennard-Jones particle system, *T*_c_ scales with *ε*. This scaling relation illustrates that phase separation is driven by attractive interactions between constituent macromolecules; the stronger the attraction, the higher the critical temperature for phase separation. An approximate expression for the interfacial tension is^26^

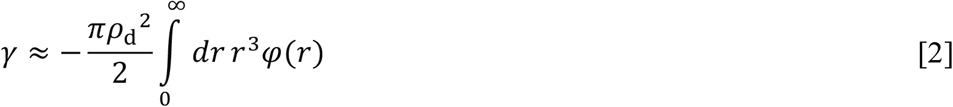

where *ρ*_d_ is the density of the dense phase. This approximation illustrates that the interfacial tension also scales with the interaction strength. Since both *T*_c_ and *γ* scale with *ε,* we can expect correlation between *T*_c_ and *γ* when intermolecular interactions are perturbed. Our results below indeed conform to this expectation. We also present unexpected behaviors, indicated by anomalous *γ* for certain mixtures.

## RESULTS AND DISCUSSION

We studied binary mixtures of Lennard-Jones particles or chains comprising 10 such particles. The strength, *ε*_DD_, of the self-interaction of driver particles is assigned a value of 1 to set the energy scale. The counterpart, *ε*_RR_, for regulator particles has a value of 0.9. Similar to previous work,^15, 18^ we study three different values for *ε*_DR_, i.e., 0.8, 1.0, and 1.2. The first *ε*_DR_ value is less than *ε*_RR_ whereas the third *ε*_DR_ value is greater than *ε*_DD_. For a given *ε*_DR_ value, we also study the whole range of mixing ratios between driver and regulator particles or chains, with regulator mole fraction, *x*_R_, at 0 (pure driver), 0.2, 0.4, 0.6, 0.8, and 1 (pure regulator). Lastly, for each *ε*_DR_and *x*_R_, we study a range of temperatures, for particles starting at 0.65 and going all the way toward *T*_c_with increments of 0.02 whereas for chains starting at 1.7 and going all the way toward *T*_c_ with increments of 0.1. We use the term “corresponding” to refer to a particle system and a chain system that have the same values for *ε*_DR_ and *x*_R_.

### Strength of D-R Interactions Dictates Regulatory Effects on Critical Temperature

From the MD simulations, we obtain the total densities, *ρ*_b_ and *ρ*_d_, of the two components in the bulk and dense phases (see Computational Methods and Figure S1a) at a series of temperatures (Figure 1a for particles and Figure 1b for chains). We fit the resulting binodals to the following equations:

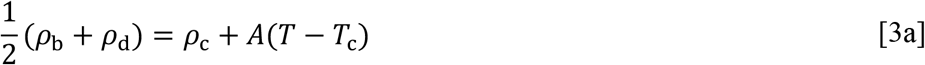

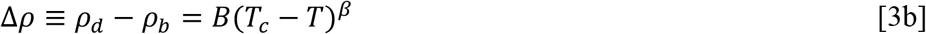

where *ρ*_c_ is the critical density, *A* and *B* are constants, and the exponent *β* is set to 0.32. Eq [3a] is called the law of rectilinear diameters,^27^ whereas eq [3b] is a scaling relation.^27-28^

**Figure 1.**
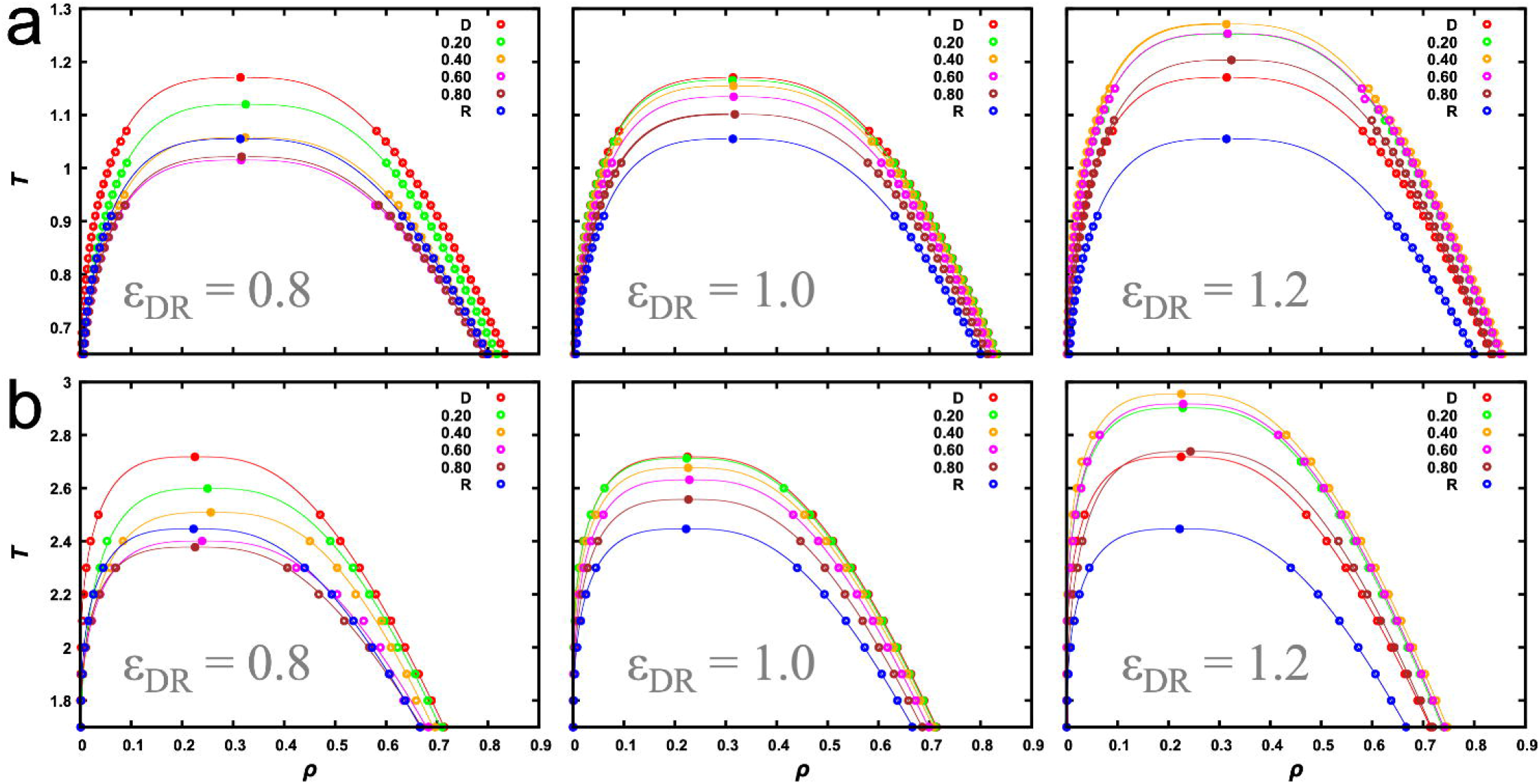
Binodals of (a) particle and (b) chain systems. The driver-regulator interaction strength, *ε*_DR_, is listed near the bottom of each panel; the mole fraction of the regulator species is listed at the top right. Symbols are densities of the two phases at a given temperature, determined by fitting the density profile to a hyperbolic tangent function. The curves are fits of the binodals to eq [3]. Given the greater uncertainties near the critical region, we successively removed data closest to *T*_c_, until the fitted values of *T*_c_ and *ρ*_c_ were stable.^15^

The critical temperature for the pure driver particle system is 1.171, in good agreement with a value 1.16 in a previous study for the same cutoff.^12^ The pure regulator particle system is identical to the pure driver counterpart, except that the interaction energy is reduced by a factor of 0.9. Correspondingly the critical temperature is expected to be reduced to 1.171 × 0.9 = 1.054, which is nearly identical to the *T*_c_ value, 1.055, determined from our simulations of the pure regulator particle system. When regulator particles are mixed with driver particles, very different changes n *T*_c_ are observed depending on the value of *ε*_DR_ relative to *ε*_DD_(Figures 1a and 2a). At *ε*_DR_ = 0.8, as *x*_R_ increases from 0, *T*_c_ gets lower. Then at *x*_R_ = 0.6 and 0.8, where the regulator becomes the major species, *T*_c_is even lower than that for pure regulator particles. Of course as *x*_R_ → 1, *T*_c_ must come back to the value for pure regulator particles. The dependence of *T*_c_ on *x*_R_ for at *ε*_DR_ = 0.8 has the shape of an upward parabola.

At *ε*_DR_ = 1.2, *T*_c_initially increases with increasing *x*_R_, which is just opposite to the situation at *ε*_DR_= 0.8. At *x*_R_ = 0.6 the trend is reversed and *T*_c_ decreases all the way till *x*_R_ = 1, but still is above that for pure driver particles even at *x*_R_ = 0.8. The overall shape of the dependence of *T*_c_ on *x*_R_ is a downward parabola. At *ε*_DR_ = 1.0, the dependence of *T*_c_ on *x*_R_ is close to a linear interpolation between the two end values (i.e., for pure driver and pure regulator), but with a small downward curvature.

The *T*_c_ trends at the different *ε*_DR_ values are qualitatively in line with our conclusion in previous studies,^15,18^ in which regulators with *ε*_DR_ < *ε*_DD_ and with *ε*_DR_ > *ε*_DD_ are classified as weak-attraction suppressors and strong-attraction promotors, respectively. Weak-attraction regulators suppress phase separation by partitioning into the dense phase and thereby replacing some of the stronger driver-driver interactions with weaker driver-regulator interactions; strong-attraction regulators do exactly the opposite at low *x*_R._ One difference between the present work and our previous studies lies in the self-interaction of regulator particles. In the previous studies, regulator particles experience only steric repulsion among themselves, whereas here they experience Lennard-Jones interactions, albeit with a weaker strength than that for the driver particles. Therefore, in our previous studies, the extent of weak-attraction suppression of phase separation, as measured by the decrease in *T*_c_, increases monotonically with *x*_R_, whereas here the decrease in *T*_c_ reaches a maximum at an intermediate *x*_R_. Similarly, in our previous studies, once strong-attraction regulators switch from being promotors to suppressors at a certain *x*_R_, *T*_c_ decreases indefinitely with further increase in *x*_R_, but here *T*_c_ decreases at most to the value for pure regulator particles.

The *T*_c_ trends at the three *ε*_DR_ values for binary mixtures of Lennard-Jones chains (Figures 1b and 2b) qualitatively parallel those for the Lennard-Jones particles. Quantitatively, two differences are worth noting. First, the critical temperatures of the chain systems are higher, by a factor *α*_0_ ≈ 2.3, than the corresponding particle systems. Second, the binodals shift toward lower densities. These two differences in the binodals of particle and chain systems have been pointed out previously based on predictions of perturbation theories.^14^

### Particle Systems and Chain Systems Have Approximately Equal Interfacial Tensions at Matching Temperatures

For each binary mixture at a given temperature, we calculated the interfacial tension according to the Kirkwood-Buff method.^29^ We noted already that, in either the particle case or the chain case, the two pure systems (at *x*_R_ = 0 and 1) are identical except that the interaction energies are at a ratio of 0.9. As can be seen by their units (i.e., *ε*/*k*_B_ and *ε*/σ^2^, respectively), both of *T* and *γ* scale with the interaction strength. Therefore we expect that the values for the pure driver systems, (*T*_R_, *γ*_R_), should equal the counterparts for the pure regulator systems, (*T*_D_, *γ*_D_), when the latter quantities are both scaled up by a factor of 1/0.9. That is, *γ*_D_ = *γ*_R_/0.9 when *T*_D_ = *T*_R_/0.9. The results of our MD simulations indeed agree with this expectation (Figure S2).

Figure 3a displays illustrative results for a mixture, comprising Lennard-Jones particles at *ε*_DR_ = 1.2 and *x*_R_ = 0.4. In all cases, *γ* gradually decreases toward 0 as *T* increases toward *T*_c._ The dependence of *γ* on *T* fits well to the scaling relation^27-28^

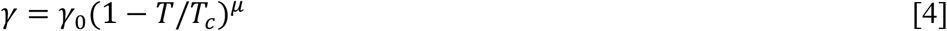

with *μ* = 1.255 (Figure 3a inset and Figure 4a, b). This fit provides a second route to the determination of *T*_c_. The resulting values are in very close agreement with those determined by fitting the binodals (Figure 3b).

**Figure 3.**
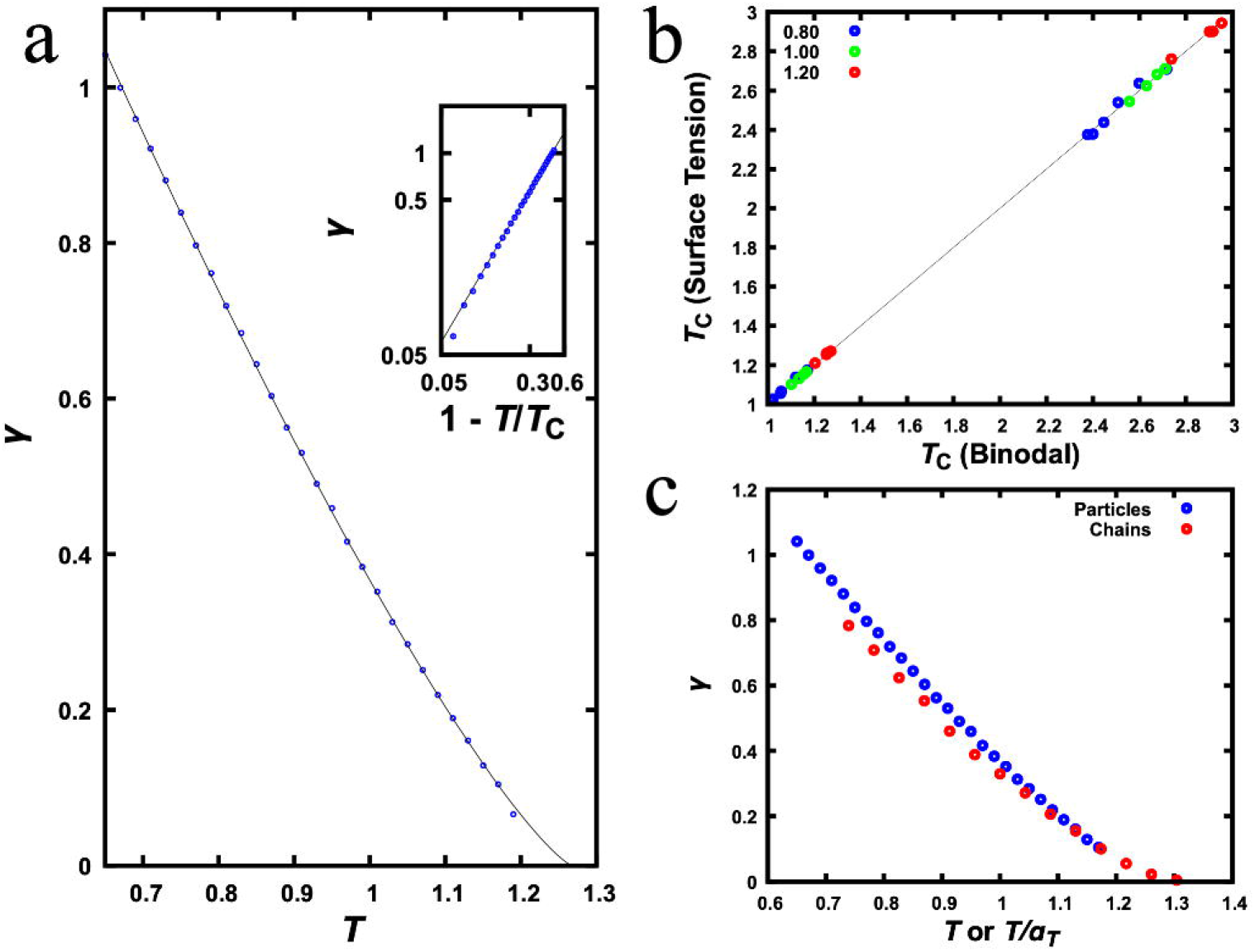
Temperature dependence of interfacial tension. (a) Results for a mixture of Lennard-Jones particles at *ε*_DR_ = 1.2 and *x*_R_ = 0.4. The curve is a fit to eq [4]. The inset displays a similar plot but with both *γ* and 1 − *T*/*T*_c_ on a logarithmic scale. (b) Comparison of critical temperatures determined via two different routes, by fitting either binodal or interfacial tension. The line represents perfect agreement between the two routes. (c) Comparison of interfacial tensions of the corresponding particle and chain systems at *ε*_DR_ = 1.2 and *x*_R_ = 0.4. For the chain system, the abscissa represents temperature scaled down by a factor *α_T_* = 2.3.

**Figure 4.**
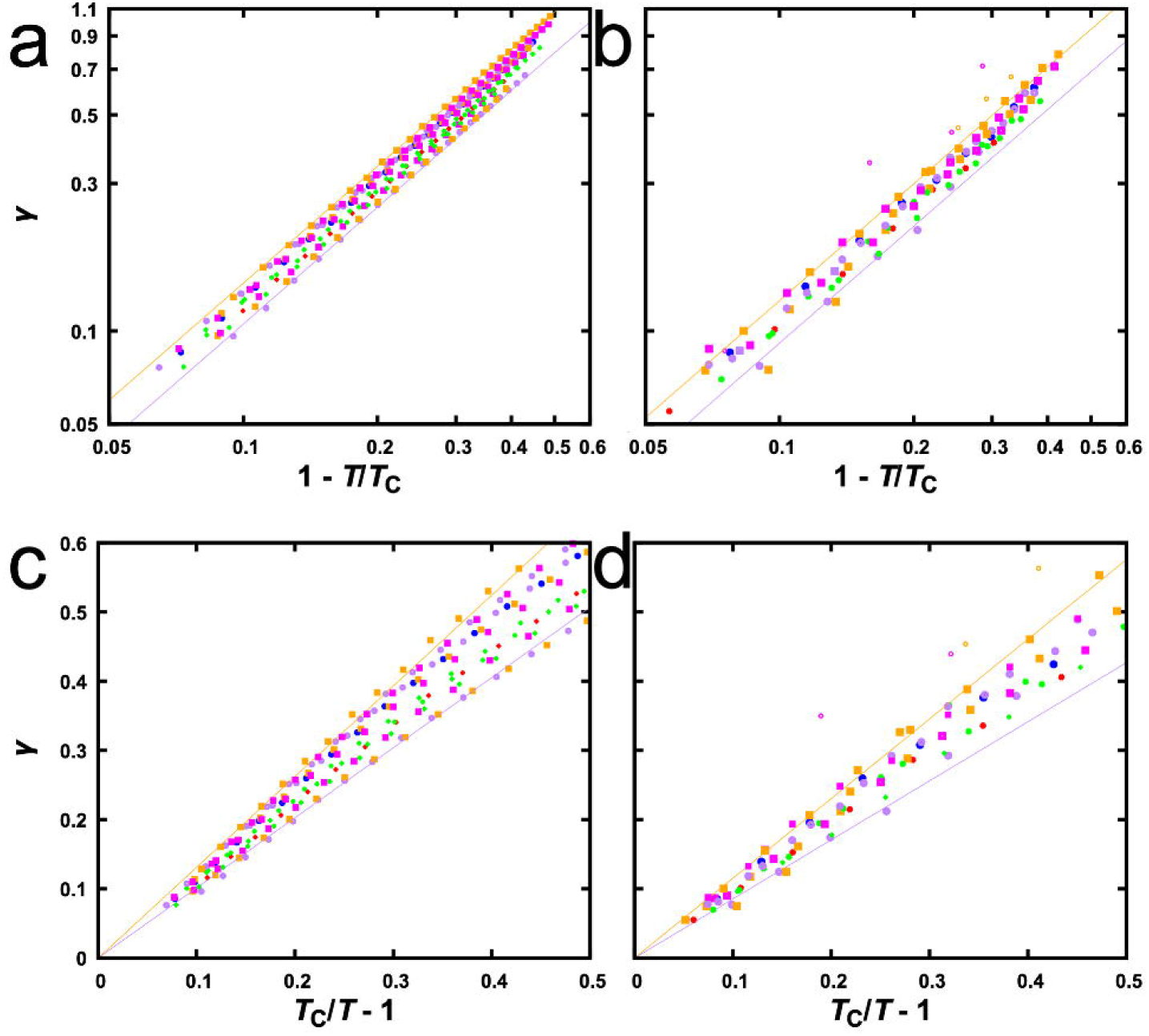
Scaling of interfacial tension with *T* or with *T*_c_. (a) Scaling of *γ* with 1 − *T*/*T*_c_ for the particle systems, displayed on a log-log scale. The fits to eq [4] for two sets of data near the borders are shown as lines. (b) Corresponding plots for the chain systems. (c) Scaling of *γ* with *T*_c_/*T* − 1 for the particle systems. Fits to eq [7], with *T*_c_ fixed at the values from fitting interfacial tensions, for two sets of data (each for a given mixture in a range of *T*) near the borders are shown as lines. (d) Corresponding plots for the chain systems. Anomalous *γ* values due to multiphase coexistence are shown as open symbols in panels (b) and (d).

Comparing the interfacial tensions of the particle systems and the corresponding chain systems, we find that they are approximately equal at matching temperatures, especially at *T* close to *T*_c_ (Figure 3c). By matching temperatures, we mean that a temperature *T* for a particle system is equivalent to a temperature *α_T_T* for the corresponding chain system. The similarity in interfacial tension between corresponding particle and chain systems will be further discussed below.

Combining eqs [3b] and [4], we obtain a third scaling relation

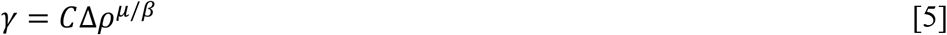

The exponent is expected to be 3.92, based on the above *β* and *μ* values. Our data fit well to this scaling relation, with exponents very close to the expected value (mean and standard deviation of exponents at 3.93 ± 0.07; Figure S3). The last scaling relation indicates that, other things being equal, *γ* grows with increasing difference in density between the dense and bulk phases. This density difference and hence *γ* decrease toward 0 as *T* approaches *T*_c_.

### Regulatory Effects on *T*_c_ and on *γ* Match Each Other

At a given temperature, the overall mole fraction, *x*_R_, of the regulator species affects the compositions of the two coexisting phases (Figure 1) and therefore the interfacial tension. The dependence of *γ* on *x*_R_ for Lennard-Jones particles was reported by Stephan et al.^24^ at a single temperature. Here we studied the interfacial tensions of both particle and chain systems over a range of temperatures. The results are illustrated in Figure 2 for the particle systems at *T* = 0.95 and the chain systems at *T* = 2.2. Interestingly, the effects of regulator particles and chains on *γ* at a fixed *T* match almost perfectly with the corresponding effects on *T*_c_. Simple reasoning as presented in the Introduction gave us an inkling for a correlation between *γ* and *T*_c_, but the degree of correlation exhibited by the data is somewhat surprising.

**Figure 2.**
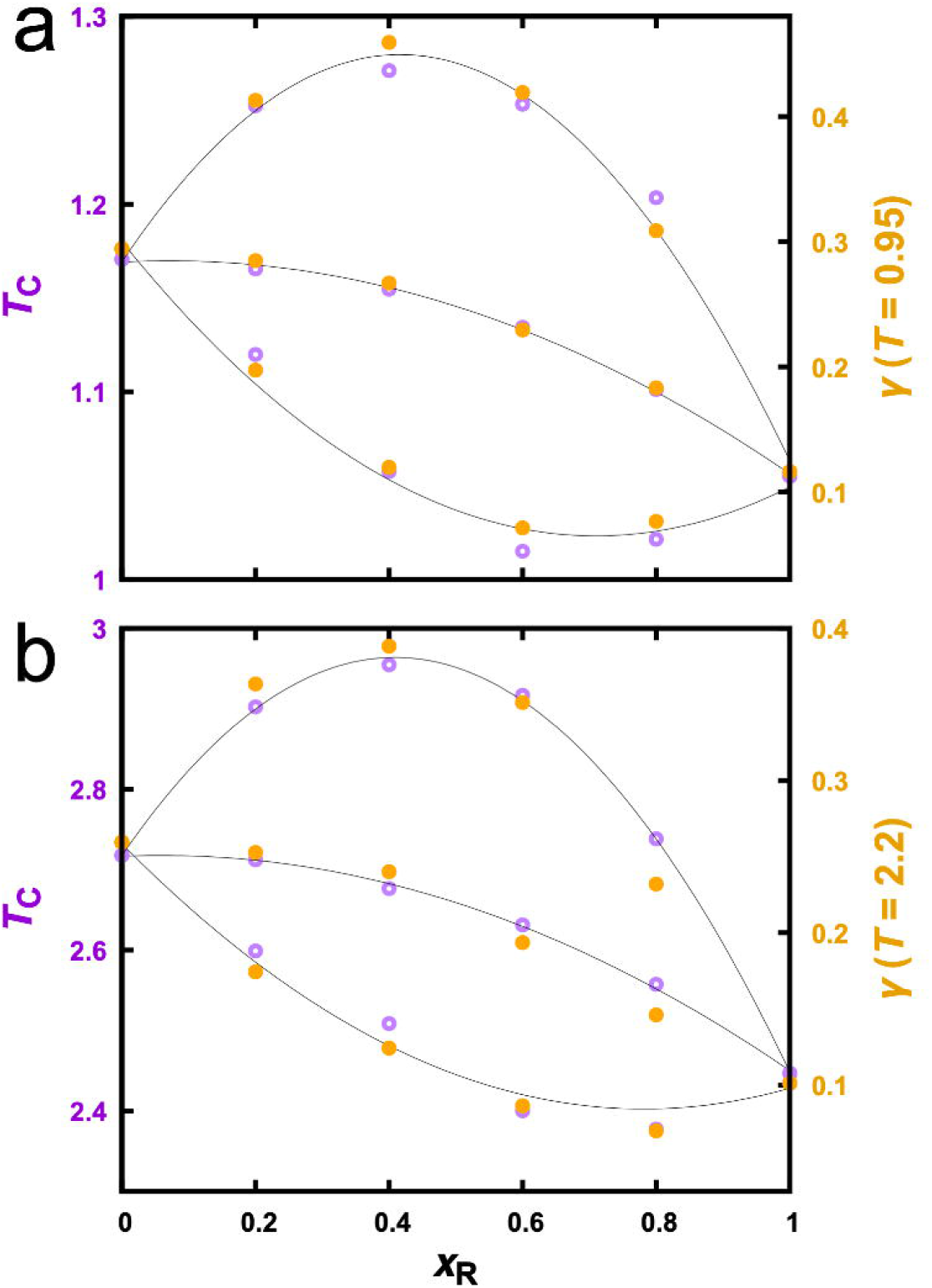
Regulatory effects on *T*_c_ and on *γ*. (a) Particle systems. (b) Chain systems. *T*_c_ values are determined by fitting binodals. Curves are parabolic fits to guide the eye. Data at *ε*_DR_ = 1.2, 1.0, and 0.8 are displayed from top to bottom in each panel.

Given the high degree of correlation between *T*. and *γ*, the trends described above for *T*_c_ at different *ε*_DR_ also translate into those for *γ*. That is, at *ε*_DR_ = 0.8, regulators decrease *γ*, even to values below *γ*_R_ when *x*_R_ is at 0.6 and 0.8; at *ε*_DR_ = 1.2, regulators initially increase *γ* but then turn over at *x*_R_ = 0.6, but *γ* is still higher than *γ*_D_ even at *x*_R_ = 0.8; at *ε*_DR_ = 1.0, the regulatory effects on *γ* are close to a linear interpolation between *γ*_D_ and *γ*_R_ but with a small downward curvature.

We find that the match in the regulatory effects on *T*_c_ and on *γ* is better when *T* is closer to *T*_c._ While Figure 2a compares *T*_c_ with *γ* at *T* = 0.95 for particle mixtures, when a similar plot is made for *γ* at *T* = 0.65 we see larger deviations of the *γ* trends from the *T*_c_ trends (Figure S4a). For the chain mixtures at *ε*_DR_ = 0.8, we even see *γ* values at *x*_R_ = 0.4 and 0.6 that buck the trend set by *T*_c_ (Figure S4b; see also Figure S3b). We will elaborate below on this anomalous behavior.

### *γ* and *T*_c_ Have a Linear Relation

The foregoing results show that regulatory effects on *T*_c_ can predict the corresponding effects on *γ* at any temperature. Figure 2 even suggests a linear relation between *γ* and *T*_c_. To provide a justification, let us manipulate the scaling relation given by eq [4] (see Figure 4a, b), for *T* close to *T*_c_. Let *δ* = *T*_c_ − *T*. Expressing *T*_c_ in terms of *δ* and carrying out a Taylor expansion, we obtain

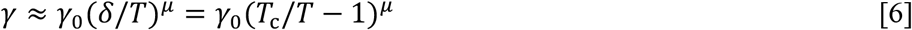

This expression suggests that we plot *γ* against *T*_c_/*T* − 1 and fit the resulting relation to a power law. Such fits yield exponents that are very close to 1, or an approximate linear relation between *γ* and *T*_c_,

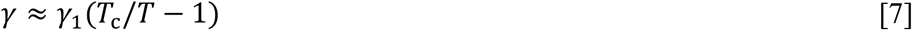

Figure 4c, d display this linear relation, for *T*_c_/*T* − 1 < 0.5. For the particle systems, the slope *γ*_1_ falls within the range of 1.01 to 1.31; the corresponding range for the chain systems is 0.85 to 1.15.

The similar slopes of the particle and chain systems are supported by the finding made clear above by Figure 3c, which is that corresponding particle and chain systems have nearly equal interfacial tensions at matching temperatures close to their respective *T*_c_ values. When matching temperatures, we scale both *T*_c_ and *T* by the same factor *α*_0_, thus leaving *T*_c_/*T* the same for the particle and chain systems. If both *γ* and *T*_c_/*T* − 1 are the same, then the slope *γ*_1_ has to be the same for corresponding particle and chain systems.

A linear relation, or indeed any one-to-one relation, between *T*_c_ and *γ* is highly significant. It means that, when a condensate-forming system experiences a change in macromolecular composition or a change in intermolecular interactions (e.g., by mutation or posttranslational modification, or by a perturbation in solvent conditions such as temperature, pH, or salt), the resulting change in *T*_c_ can be used to predict the change in *γ* and vice versa.

### Disparity in Self- and Cross-Species Interactions Drives Multiphase Coexistence

Finally let us focus our attention to the anomalous interfacial tension observed on the chain mixtures at *ε*_DR_ = 0.8, *x*_R_ = 0.4 and 0.6, and *T* below 2.2 (Figures 4b, d; S3b; and S4b). Inspecting snapshots from the simulations, we were surprised to discover that the dense phase is not a homogeneous mixture of the two components (Figure 5a). Instead, driver chains and regulator chains demix to form two distinct dense phases: a driver-rich slab at the center, bordered by two regulator-rich slabs on the two sides (Figure 5b). This type of multiphase coexistence has been reported in many experimental studies^1, 7, 9-10^ and in some recent computational studies,^10, 30-31^ and may underlie the organization of many membraneless organelles.^1,4-6, 8^ It is interesting that our highly simplified model systems recapitulate this complex phenomenon, affording us an opportunity to elucidate its general physical basis.

**Figure 5.**
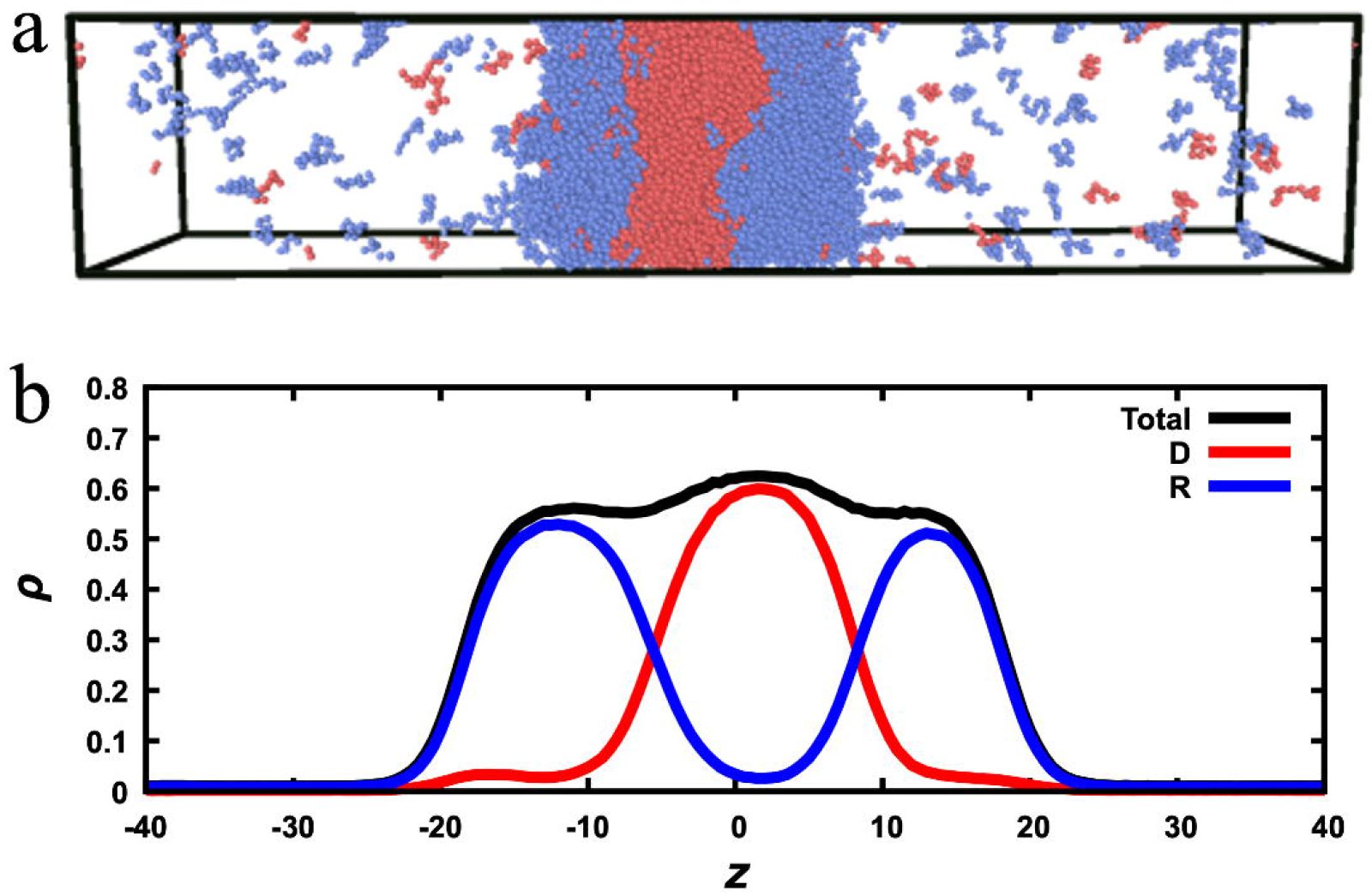
Multiphase coexistence observed for the chain systems. (a) A snapshot of the system at *ε*_DR_ = 0.8 and *x*_R_ = 0.4. Driver and regulator chains are in red and blue, respectively. (b) Density profiles for the driver species, regulator species, and both species combined.

Our initial results here already allow us to draw some tentative conclusions. First, in the ranges of parameters studied, we observed driver-regulator demixing only at low *ε*_DR_ and low *T*. Demixing means that driver chains want to stay with driver chains and regulator chains want to stay with regulator chains. A *ε*_DR_ that is lower than both *ε*_DD_ and *ε*_RR_ would explain why that happens, as doing would minimize the less favorable cross-species interactions and maximize the more favorable self-interactions of each species. Low *T* would accentuate the disparity between the cross-species interaction strength *ε*_DR_ and the self-interaction strengths *ε*_DD_ and *ε*_RR_, as the contrast between the corresponding Boltzmann factors would be magnified at low *T*. Second, demixing was observed only at *x*_R_ = 0.4 and 0.6, namely close to a 1:1 ratio between the driver and regulator species. The 1:1 molar ratio is where the two species would have the highest chance of interacting with each other if they were homogeneously mixed. That demixing was observed only around this molar ratio directly indicates that demixing results from the avoidance of cross-species interactions. A full investigation into the physical basis of multiphase coexistence will be carried out in the future.

Multiphase coexistence creates problems for the analysis methods that are designed for two-phase systems. For example, the profiles of the component densities or even the total density are no longer a single transition as a function of *z* (compare Figure S1a and Figure S1b). There is an additional mini-transition between the driver-rich phase and the regulator-rich phase. When we fit the profile of the total density to a hyperbolic tangent function, the high-density plateau represents the average density of the two dense phases. Likewise, instead of the single type of interface (i.e., between the dense and bulk phases), we now have two types of interfaces, one between the driver-rich phase and the regulator-rich phase, and one between the regulator-rich phase and the bulk phase. The interfacial tension determined by the Kirkwood-Buff method is probably some kind of average of the surface tensions at these interfaces, but the exact nature remains to be clarified.

## CONCLUSION

The results for both mixtures of Lennard-Jones particles and mixtures of Lennard-Jones chains have demonstrated that, when intermolecular interactions are perturbed (by, in particular, a compositional variation), the resulting changes in critical temperature and in interfacial tension are highly correlated. Our previous studies have shown that, depending on the relative strength *ε*_DR_/*ε*_DD_, *T*_c_ as a function of *x*_R_ follows distinct trends, and the predicted trends have been validated by experiments.^15, 18^ Now we can predict that *γ* as a function of *x*_R_ follows exactly the same trends. Extending the experimental test to the predictions on *γ* will be very interesting. Beyond this experimental test, the results obtained here deepen our general understanding of how interfacial tensions of biomolecular condensates are affected by macromolecular compositions or interactions, and therefore how biomolecular condensates mediate cellular functions. While the present work focused on changes in *T*_c_ and *γ* brought by compositional variations, the correlation between them should extend to other kinds of perturbations in intermolecular interactions, e.g., by mutation or posttranslational modification, or by a variation in solvent conditions such as temperature, pH, or salt. The close correlation between *T*_c_ and *γ* allows the data for one property to be used to predict the other property. Lastly, the observation that the highly simplified model systems studied here exhibit multiphase coexistence puts us in a position to learn what drives this phenomenon. A tentative conclusion is that the multiphase coexistence observed here is driven by disparity in self- and cross-species interactions; that is, the two species demix in order to minimize the less favorable cross-species interactions and maximize the more favorable self-interactions of each species. Much more can be learned in future studies.

## COMPUTATIONAL METHODS

### Molecular Dynamics Simulations

Binodals and interfacial tensions were calculated from molecular dynamics (MD) simulations, using the open-source software package HOOMD-blue (version 2.5.0) on graphics-processing units.^32^ We largely followed Silmore et al.^25^ but studied two sets of binary mixtures: one comprising Lennard-Jones particles and the other comprising chains of 10 Lennard-Jones particles. The two species of particles have different self-interaction strengths. The interaction strength of the “driver” species, *ε*_DD_, is set to *ε*, which is the unit of energies. Several values of the cross-species interaction strength, *ε*_DR_, are studied. The “regulator” species has an interaction strength of *ε*_RR_ = 0.9*ε*. Driver chains are homopolymers of driver particles whereas regulator chains are homopolymers of regulator particles. All the particles have the same diameter σ, which is the unit of lengths, and the same mass *m*. The units for number density, temperature, interfacial tension, and time are σ^-3^, *ε*/*k*_B_, *ε*/σ^2^, and 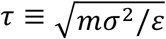, respectively.

Periodic boundary conditions were applied in the simulations. In the unit cell, the total number of particles was 20000 for the particle systems and correspondingly the total number of chains were 2000 for all the chain systems. The mole fraction, *x*_R_, of the regulator species was varied to span the entire range, from pure drivers (*x*_R_ = 0) to pure regulators (*x*_R_ = 1). The Lennard-Jones interactions were truncated and shifted, with a cutoff 3σ for all the particle systems and 6σ for all the chain systems. For chains, Lennard-Jones interactions were not applied between neighboring particles. Instead, they were constrained by a harmonic potential with a spring constant 75,000*ε*/σ^2^ and an equilibrium length σ. In addition to volume, the temperature of the system was also kept constant by running Langevin dynamics with a friction coefficient of 0.1*m*/τ. The integration timestep was 0.005τ for the particle systems and 0.001τ for the chain systems.

The procedure to prepare the system for phase separation is illustrated in Figure 6. To start, particles or chains of particles were randomly placed in a cubic box with dimensions *L_x_* = *L_y_* = *L_z_* = 60σ (corresponding to a low initial density of 0.093; Figure 6a), and energy minimized to relieve clashes. The particles were then linearly compressed at a high temperature of *T* = 4*ε*/*k*_B_ for 5000 timesteps, to reduce the cubic box to half of its original dimension in each direction (density at 0.74; Figure 6b). Finally, the box, but not the particles, was elongated in the *z* direction to *L_Z_* = 150σ, creating empty space on both sides of the compressed particles (Figure 6c). Hereafter, we set the temperature to a desired value (below *T*_c_) and started the simulation to allow the system to relax and reach a phase-separated equilibrium (Figure 6d). The total simulation length was 10 million steps; the second 5 million steps were used for calculating average properties, as described next.

**Figure 6.**
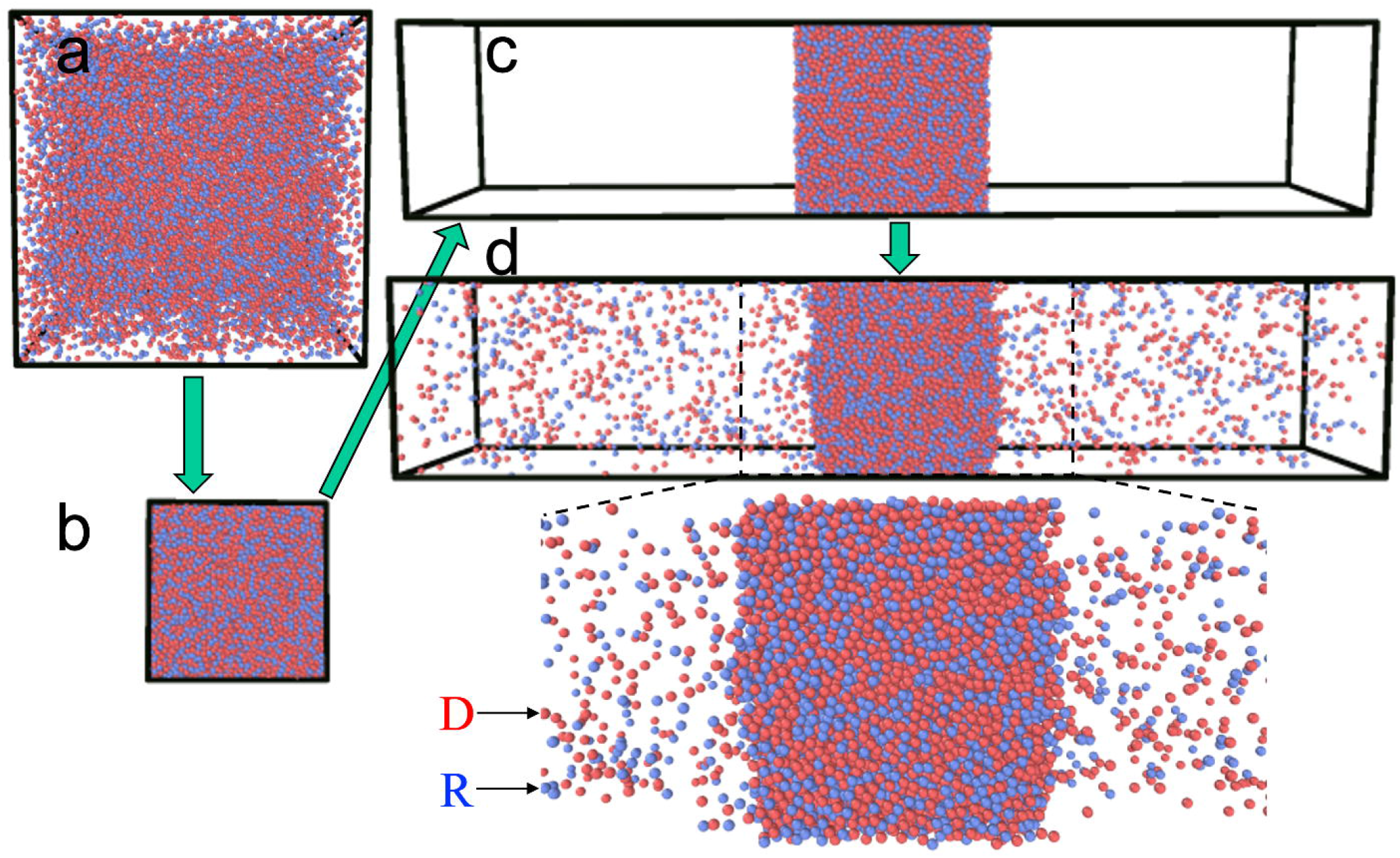
Setup and simulation to reach phase-separated equilibrium. (a) System in an initial cubic box. (b) System after 2-fold compression in each direction. (c) Enlarged box in *z* direction. (d) Two-phase equilibrium. An enlarged view of the dense phase and the neighboring regions in the bulk phase is shown below.

### Determination of Densities in the Dense and Bulk Phases

500 snapshots in the equilibrated portion of the simulation were saved (at intervals of 10000 timesteps). The density profile, *ρ*(*z*), along the *z* direction was calculated by dividing the simulation box of each snapshot into slabs of thickness 0.5σ along *z*. The total number of particles (or particles of a given species) in each slab divided by the slab volume gave an estimate for the density at that particular *z*. This estimate was then averaged over the 500 snapshots to yield *ρ*(*z*).

To obtain the densities, *ρ*_d_ and *ρ*_b_, in the dense and bulk phases, we fitted the density profile for *z* between 0 and 75σ (covering half of the simulation box) to the following hyperbolic tangent function

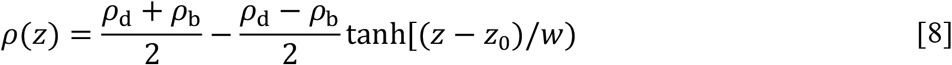

where *z*_0_ represents the midpoint of the interface between the two phases, and *w* is a measure of the width of the interface. As illustrated in Figure S1a, the fitting works very well. The corner leading to the *ρ*_b_ plateau given by the tanh function can be a bit too gradual, but that does not affect the accuracy of the value determined for *ρ*_b_.

### Determination of Interfacial Tension

The interfacial tension was determined according to the Kirkwood-Buff method,^29^ in which *γ* is expressed as

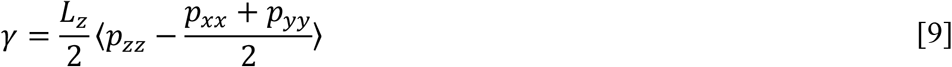

where *p_xx_, p_yy_*, and *p_yy_* are the diagonal elements of the pressure tensor, and the brackets indicate an equilibrium average. We calculated these diagonal elements on snapshots separated by 10 timesteps and averaged them over the 0.5 million such snapshots in the equilibrated portion of the simulation.

## Supplementary Material

Four additional figures (Figures S1 – S4) showing the fit of density profiles to a hyperbolic tangent function; the match in interfacial tension between pure driver systems and pure regulator systems after scaling; the scaling relation between interfacial tension and density difference between the two phases; and interfacial tensions at low temperatures.

## ACKNOWLEDGMENT

This work was supported by National Institutes of Health Grant GM118091.

## AUTHOR CONTRIBUTION

Konstantinos Mazarakos: conceptualization, investigation, methodology. Huan-Xiang Zhou: conceptualization, supervision, writing.

