## Supplementary material for "Macromolecular Regulators Have Matching Effects on the Phase Equilibrium and Interfacial Tension of Biomolecular Condensates": Figures S1 - S4

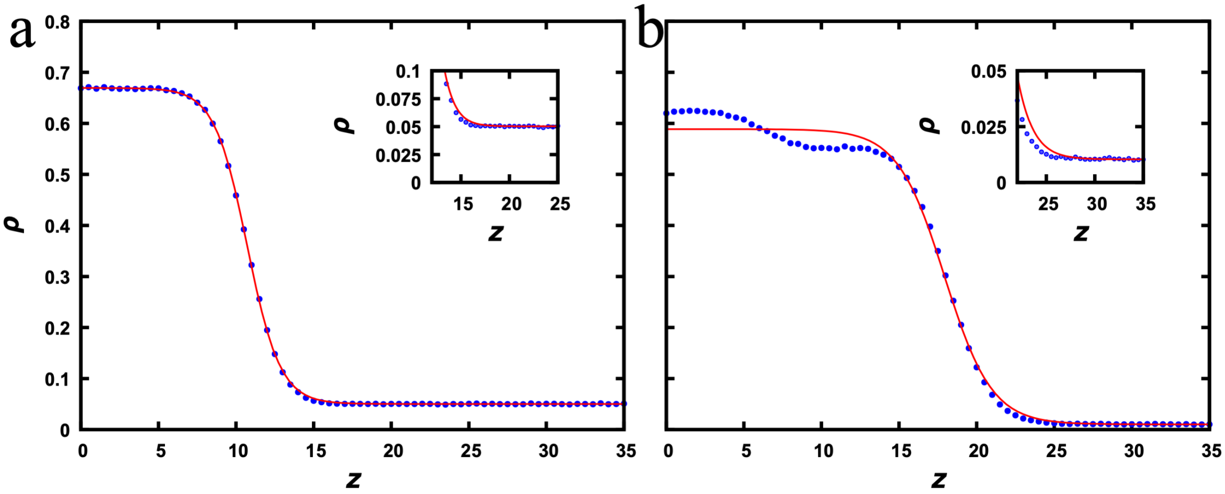


**Figure S1.** Fitting of density profiles to a hyperbolic tangent function. (a)Mixture of Lennard-Jones particles at $\varepsilon_{\mathrm{DR}} = 0.8$, $x_{\mathrm{R}} = 0.4$, and $T = 0.87$. (b) Mixture of Lennard-Jones chains at $\varepsilon_{\mathrm{DR}} = 0.8$, $x_{\mathrm{R}} = 0.4$, and $T = 2.0$. Insets display zoom into the low density region.

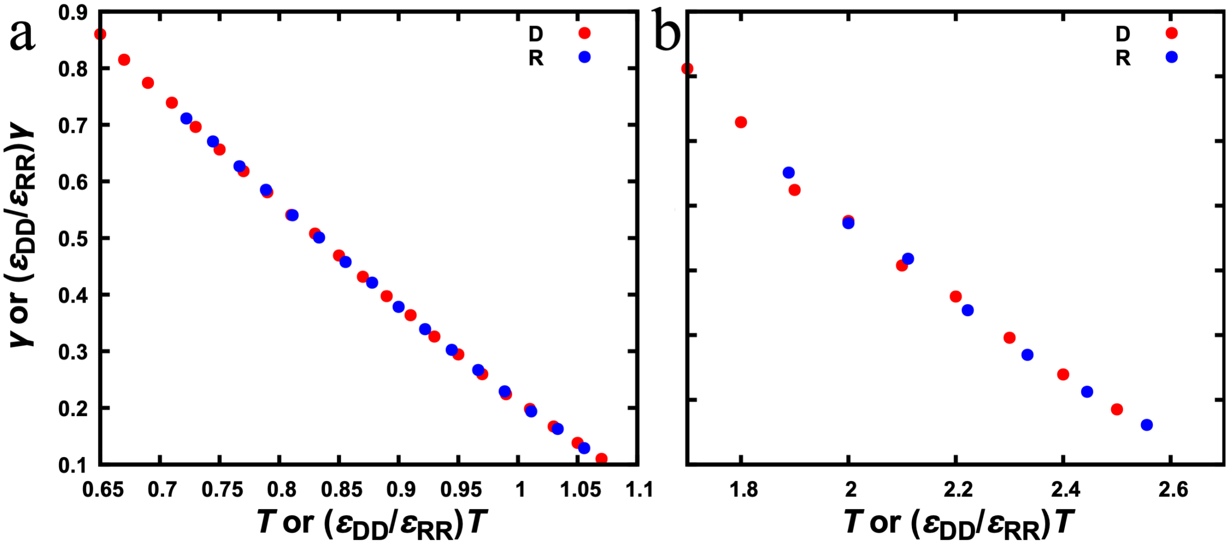


**Figure S2.** Comparison of interfacial tensions determined for the two pure systems. (a) Pure particle systems. (b) Pure chain systems. The interfacial tensions and temperatures of the pure regulator systems are scaled by the factor $\varepsilon_{\mathrm{DD}}/\varepsilon_{\mathrm{RR}} = 1/0.9$.

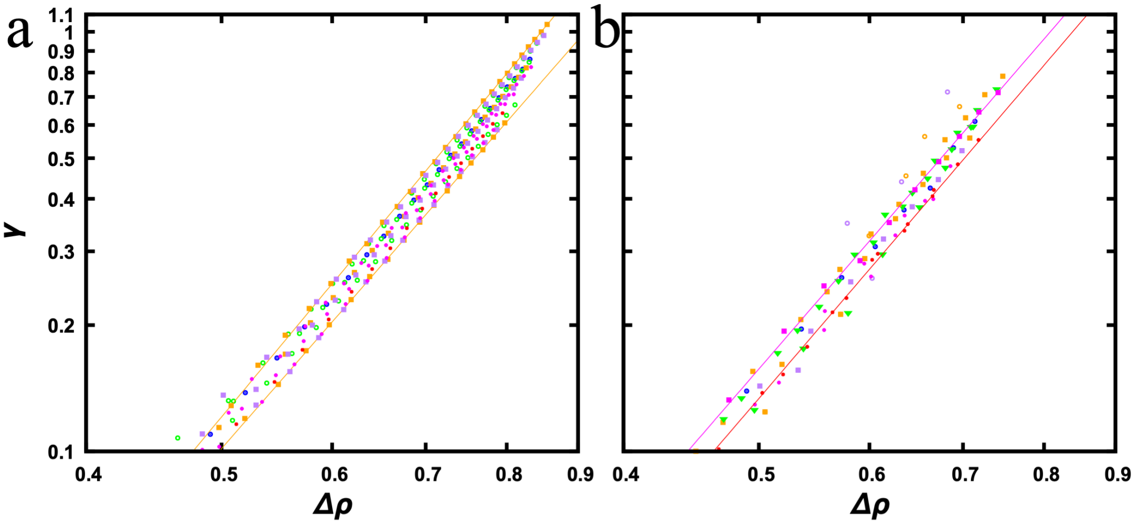


c

| | Particles | | | Chains | | |
|---|---|---|---|---|---|---|
| $\varepsilon_{\mathrm{DR}}$ \ $x$ | 0.8 | 1.0 | 1.2 | 0.8 | 1.0 | 1.2 |
| 0 | 3.98 | | | 3.98 | | |
| 0.2 | 3.70 | 3.97 | 3.95 | 3.70 | 3.97 | 3.95 |
| 0.4 | 3.81 | 3.97 | 3.99 | 3.81 | 3.97 | 3.99 |
| 0.6 | 3.89 | 3.96 | 3.93 | 3.89 | 3.96 | 3.93 |
| 0.8 | 3.93 | 3.93 | 3.83 | 3.93 | 3.93 | 3.83 |
| 1 | 3.96 | | | 3.96 | | |
| | 3.93 ± 0.07 | | | 3.93 ± 0.07 | | |

**Figure S3.** Scaling relation between interfacial tension and density difference between dense and bulk phases. (a) Particle systems. (b) Chain systems. Anomalous $\gamma$ values due to multiphase coexistence are shown as open symbols. (c) Exponents from fitting the dependence of $\gamma$ on $\Delta\rho$ to eq [7].

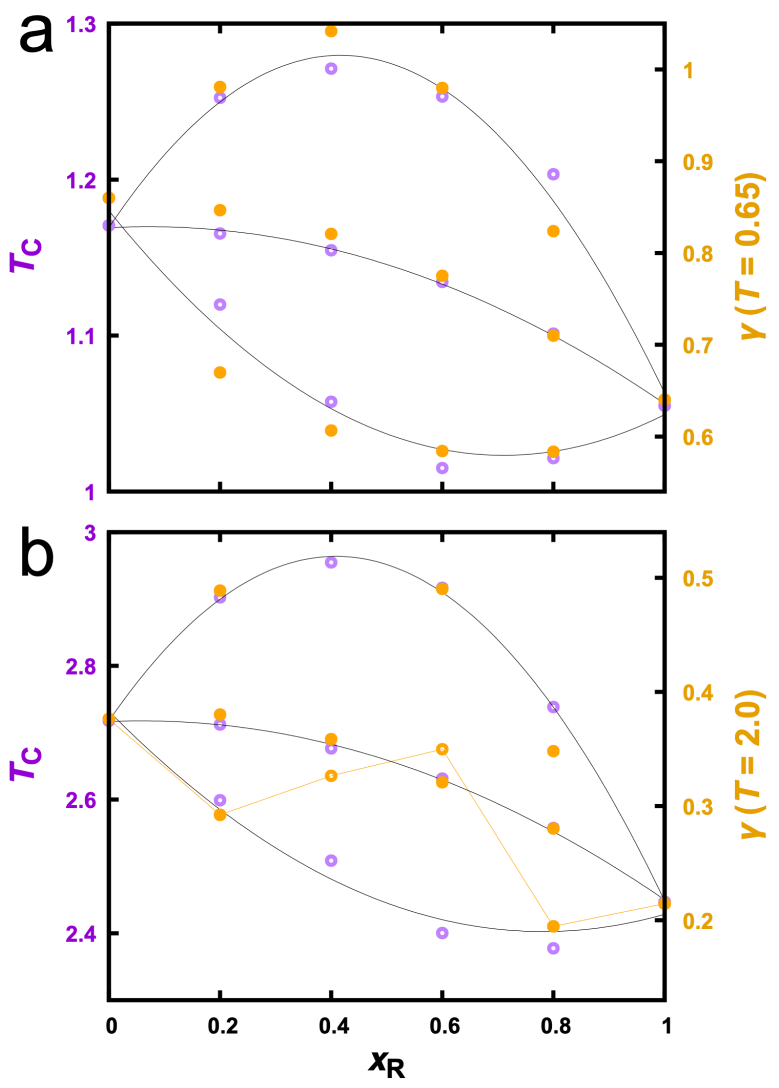


**Figure S4.** Regulatory effects on $T_c$ and on $\gamma$. (a) Particle systems. (b) Chain systems. $T_c$ values are determined by fitting binodals. Curves are parabolic fits to guide the eye. Data at $\varepsilon_{DR}$ = 1.2, 1.0, and 0.8 are displayed from top to bottom in each panel. This figure is similar to Figure 3, except that the data for $\gamma$ are at lower temperatures. Two anomalous $\gamma$ values in panel (b) at $\varepsilon_{DR}$ = 0.8 are shown as open orange symbols.
